# Subthreshold Voltage Analysis Demonstrates Neuronal Cell-Surface Sialic Acids Modulate Excitability and Network Integration

**DOI:** 10.1101/2020.04.07.030866

**Authors:** Rishikesh U. Kulkarni, Catherine L. Wang, Carolyn R. Bertozzi

## Abstract

All neurons are covered in a thick layer of carbohydrates called glycans. Glycans are modified during neurological processes and are thought to play a role in neuronal communication. We develop a voltage imaging platform for analyzing functional connectivity changes using simultaneous voltage recordings in small populations of neurons. We validate this platform using a culture model of development as well as with several pharmacological interventions. Using this platform, we show that ablation of SNA-binding glycans results in loss of functional connectivity in mouse hippocampal neurons, while ablation of MAL II binding glycans minimally perturbs functional connectivity. Altogether, our data reveal that subpopulations of glycans play different roles in maintenance of electrophysiology and provide a platform for modeling changes in functional connectivity with simultaneous voltage recordings.

## Introduction

Since the invention of intracellular electrophysiology, neuroscience has gained a broad understanding of the role of the plasma membrane and membrane-associated molecules in modulating neuronal electrophysiology(Petersen, 2017). Despite this, models of the neuronal membrane often ignore that the cell is coated with a complex matrix of neutral and charged carbohydrates called glycans(Cohen and Varki, 2010; Guzman-Aranguez and Argüeso, 2010; Shurer et al., 2019; Stanta et al., 2010) (**Scheme 1**). The brain, in particular, is enriched in certain glycans relative to the rest of the body, and loss of glycan-synthesizing and -cleaving enzymes are implicated in a wide range of neurological disorders(Schnaar et al., 2014; Grunewald et al., 2002). Glycosylation of proteins has been extensively studied in the context of gating ion channels(Ednie and Bennett, 2012). Sialic acid, which is typically found capping the carbohydrate portion of glycans, has been demonstrated to affect the gating properties of voltage-gated ion channels in a charge-dependent manner – in particular, desialylation of many voltage-gated ion channels results in a reduction in channel conductance(Ednie and Bennett, 2012; Schwetz et al., 2011). These studies have been made possible by use of genetic manipulations to remove *N-*glycosylation sites from proteins of interest. Lipid-conjugated glycans, or gangliosides, despite representing the vast majority of sialic acid in the brain(Klenk, 2009; Schnaar, 2004; Schnaar et al., 2014), have received relatively little attention due in part to their lack of genetic tractability. Previous approaches to studying gangliosides have centered around complete knockouts of a single type of ganglioside or enrichment of cells with a single ganglioside. These studies have implicated specific gangliosides in neurologically-important roles such as calcium conductance(Ledeen and Wu, 2002) and axonal outgrowth(Rodriguez et al., 2001). These studies have deepened our understanding of the roles of individual glycans in modulating neuronal properties, but an integrated understanding how glycans act as an ensemble to affect electrophysiology and network activity has yet to be established.

To understand the relationship between glycans and network activity, we sought to probe the contribution of cell-surface sialosides to the activity of multi-neuron networks. Prior studies of sialic acid established a potential role of extracellular sialic acid in maintaining neuronal excitability(Isaev et al., 2007) but did not differentiate between different populations of sialic acid due to the usage of a nonselective sialidase from *Arthrobacter ureafaciens*(Parker et al., 2012). Similarly, studies blocking neuronal neuraminidase activity changes long-term potentiation, but employed an inhibitor that lacked selectivity for specific neuraminidases(Savotchenko et al., 2015). We hypothesized that different subpopulations of cell-surface sialosides, such as those conjugated to gangliosides, may play a unique role in modulating neuronal electrophysiology. To interrogate this possibility, we required the ability to selectively remove sialic acid from populations of biomolecules in a linkage-dependent manner. We also required the ability to make membrane potential measurements from several neurons simultaneously to assay the effects of sialoside depletion on network activity. To do this, we chose to employ small molecule voltage imaging to monitor the membrane potential of several cells at once. Because voltage imaging is a direct measurement of membrane potential, it contains information about both spiking and subthreshold activity in the region of interest(Kulkarni et al., 2017).

In this work, we exploit the high-quality measurements of neuronal electrophysiology made possible by voltage imaging to develop an *in vitro* platform to measure changes in neuronal excitability, network synchronicity, and network integration. We then used a selective sialidase to acutely remove sialic acids from a ganglioside-enriched population of biomolecules on the cell surface. We compared the resulting changes in network electrophysiology to those induced by a non-selective sialidase. These data reveal the previously under-examined roles of different subpopulations of cell-surface sialic acids in modulating electrophysiology on a network scale.

**Scheme 1:**
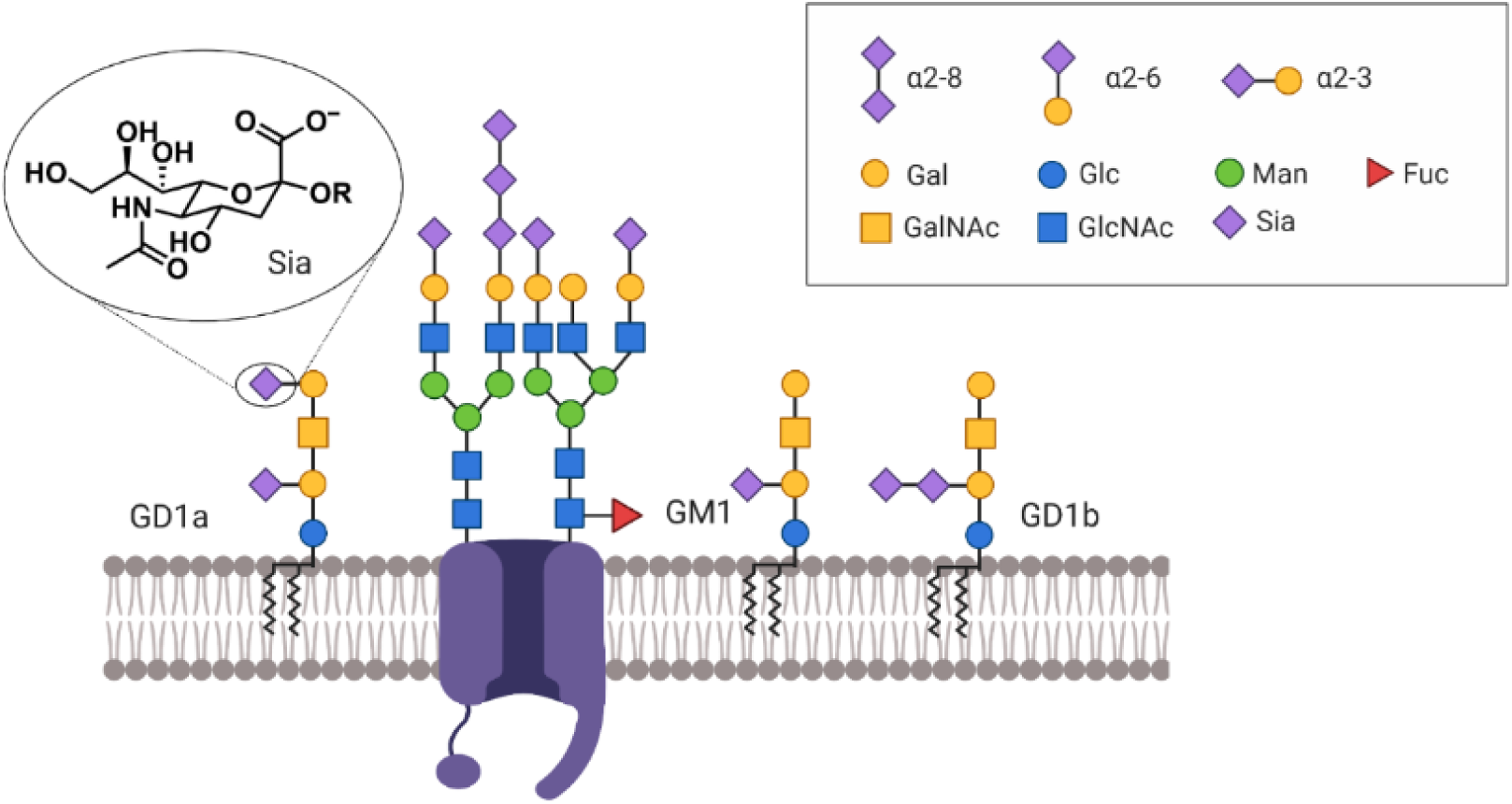
The neuronal cell membrane is densely coated with the glycans. Sialic acid (Sia) is a negatively charged residue typically found at the non-reducing end of glycans and is a significant contributor to the surface charge density of the neuronal membrane.

## Results

### *Salmonella* nanH Allows Selective Removal of 2,3-linked Sialic Acid

We hypothesize that 2,3-linked sialic acids, which are primarily found on gangliosides in the brain, play a unique role in maintaining neuronal excitability by interacting with adjacent channels and receptors in a charge-dependent manner. In order to demonstrate this, we used a pair of sialidase enzymes – *Salmonella typhimurium* nanH, which is known to selectively cleave 2,3-linked sialic acids(Gray et al., 2019; Parker et al., 2012), and *Arthrobacter ureafaciens* nanH, which promiscuously cleaves 2,3-, 2,6-, and 2,8-linked sialic acids(Parker et al., 2012). We confirmed their activity on the neuronal cell surface via incubation with each enzyme for an hour followed by staining with fluorescent dye-conjugated lectins and imaging (**Supplementary Figure 1**), which revealed that *Salmonella* nanH selectively removed 90% of the MAL II-binding sialosides (2,3-linked) with negligible activity against the SNA-binding sialosides (2,6-linked), while *Arthrobacter* nanH removed 90% of both types of visualized sialic acids (**Figure 1f, g**). With confirmation that we were able to acutely remove cell-surface sialic acid from the cell surface in a linkage-dependent or general manner, we aimed to develop a platform with which to study the effects of sialic acid on excitability and functional connectivity.

**Figure 1.**
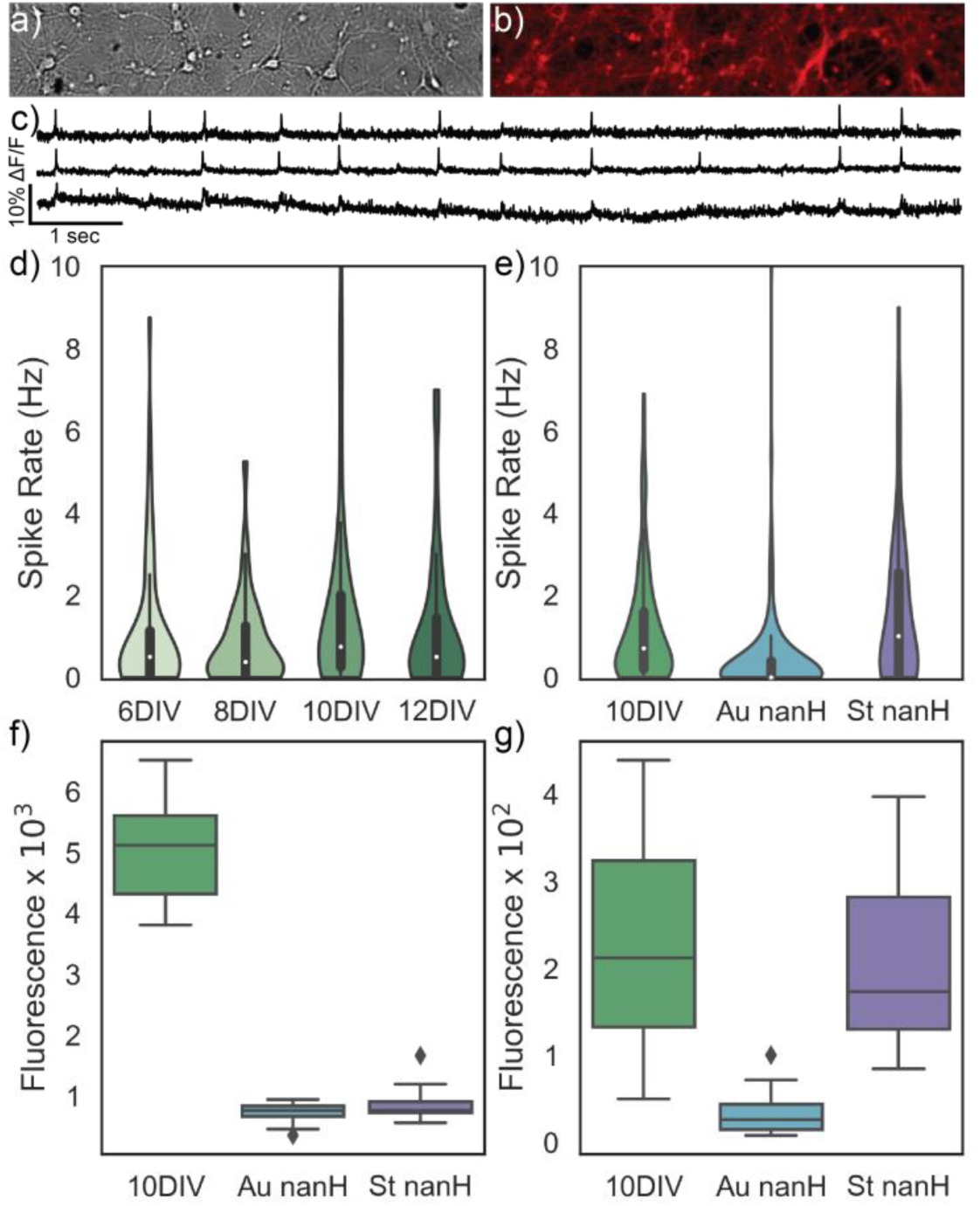
Voltage imaging reveals that sialidase treatment affects neuronal spiking rate. BeRST 1 brightly stains mouse hippocampal cultures (a, DIC image, b, 709/100 nm filter) and generates high signal-to-noise traces of neuronal membrane potential when imaged at 500 Hz (c). d) In cultured networks, average firing rate increases as the networks mature until they reach an advanced age (n = 76, 144, 191, and 125 cells across 3 animals). e) At 10DIV, treatment with *Arthrobacter* nanH results in a large reduction in firing rate, while treatment with *Salmonella* nanH results in a moderate increase in firing rate (n = 191, 108, and 98 cells per treatment across 3 animals). Staining with SNA (f) and MAL-II (g) lectins reveals that *Arthrobacter* sialidase removes 90% of both 2,3- and 2,6-linked sialic acid residues, while *Salmonella* nanH selectively removed 2,3-linked sialic acid residues (n = 17 cells per treatment). Error bars for all panels are ±STDEV.

### Sialidase Treatment Results in Changes in Neuronal Firing Rate

For voltage imaging experiments, we used the dye BeRST 1(Huang et al., 2015), a red voltage sensor from the VoltageFluor family, which has been demonstrated to directly report neuronal membrane potential with a 25 ns response time without introduction of artifacts due to dye kinetics(Beier et al., 2019). Furthermore, BeRST 1 does not affect the electrical properties of the neuronal membrane^19^. With its high signal-to-noise and fast kinetics, BeRST 1 allowed us to gather information about spiking and non-spiking behavior in several neurons simultaneously (**Figure 1a, b, c**). This enabled observation of ensembles of neurons in a network and collection of data sets of hundreds of neurons. As a baseline measure, we imaged the voltage of mouse hippocampal neurons as they matured from 6 days to 12 days in vitro (DIV). As hippocampal cultures age, they form more and more synapses, which resulted in a steady increase and plateau in firing rate (1.0 Hz, 0.8 Hz, 1.5 Hz, and 1.2 Hz, respectively) from during the imaging period until the cultures grew unhealthy due to age (**Figure 1d**). For further experiments, we used 10DIV cultures as the comparison for the administered interventions.

We then sought to remove sialic acids from the surface of these neurons to study their effects on electrophysiology. We hypothesized that *Arthrobacter* nanH treatment would result in a net decrease in spiking activity, as has been previously reported(Isaev et al., 2007). Upon treatment with *Arthrobacter* nanH, 10DIV mouse hippocampal neurons indeed demonstrated a large decrease (−0.76 Hz, p < 0.00001, Permutation test, 100,000 shuffles) in mean firing rate and population of firing cells (**Figure 1e**), in good agreement with prior studies of mammalian neuraminidase enzymes on cultured neurons. To verify that non-firing neurons were still alive, their ability to fire action potentials was verified via field stimulation. We hypothesized that *Salmonella* nanH treatment, however, which only removes 2,3-linked sialic acids, would illuminate any differences in the roles of 2,3- and 2,6-sialic acids in regulating neuronal electrophysiology. *Salmonella* nanH treatment resulted in a surprising increase (+0.42 Hz, p = 0.01254, Permutation test, 100,000 shuffles) in mean firing rate and firing population (**Figure 1e**). To elucidate the underlying mechanisms of these effects, we asked whether the changes in observed firing rates were related to changes in neuronal excitability.

### Voltage Imaging Traces Reveal Both Spiking and Subthreshold Activity

Unlike multielectrode arrays, voltage imaging enables multiplexed imaging of both spiking and subthreshold neuronal electrophysiology(Kulkarni et al., 2017). These two pieces of information can be used to infer properties of the neuronal network that would be difficult or impossible to ascertain with only spike-based metrics, such as excitability and connectivity(Adam et al., 2019; Domnisoru et al., 2013; Lampl et al., 1999; Xie et al., 2020). We developed a custom software library to process and analyze these data – in brief, the subthreshold activity is fit using an asymmetric least squares regression(Peng et al., 2010), which considers spiking data points as outliers (**Figure 2a-c**). The raw trace is then divided by the subthreshold fit, providing a flat trace that contains baseline noise and spiking time points (**Figure 2d**). A set of baseline fits (**Figure 2e**) and spike trains (**Figure 2f**) is then generated by selecting all local maxima (within 8 milliseconds) that are 4.5 standard deviations greater than the standard deviation of the baseline noise. In order to confirm that the subthreshold fit truly reflected subthreshold activity and was not an artifact of the dye, we imaged neurons in the presence of APV and NBQX, which completely abolished the variance in the subthreshold fit (**Figure 2e, Supplementary Figure 2**). Furthermore, electrically-induced action potentials in the +APV, +NBQX neurons resulted in a flat subthreshold fit, indicating that subthreshold variation was not due to an artifact resulting from interactions between the dye and the neuron (**Supplementary Figure 3**). With the “input” and “output” of each neuron now separated, we sought to determine if the changes in firing rate observed in nanH-treated neurons could be explained by changes in excitability.

**Figure 2.**
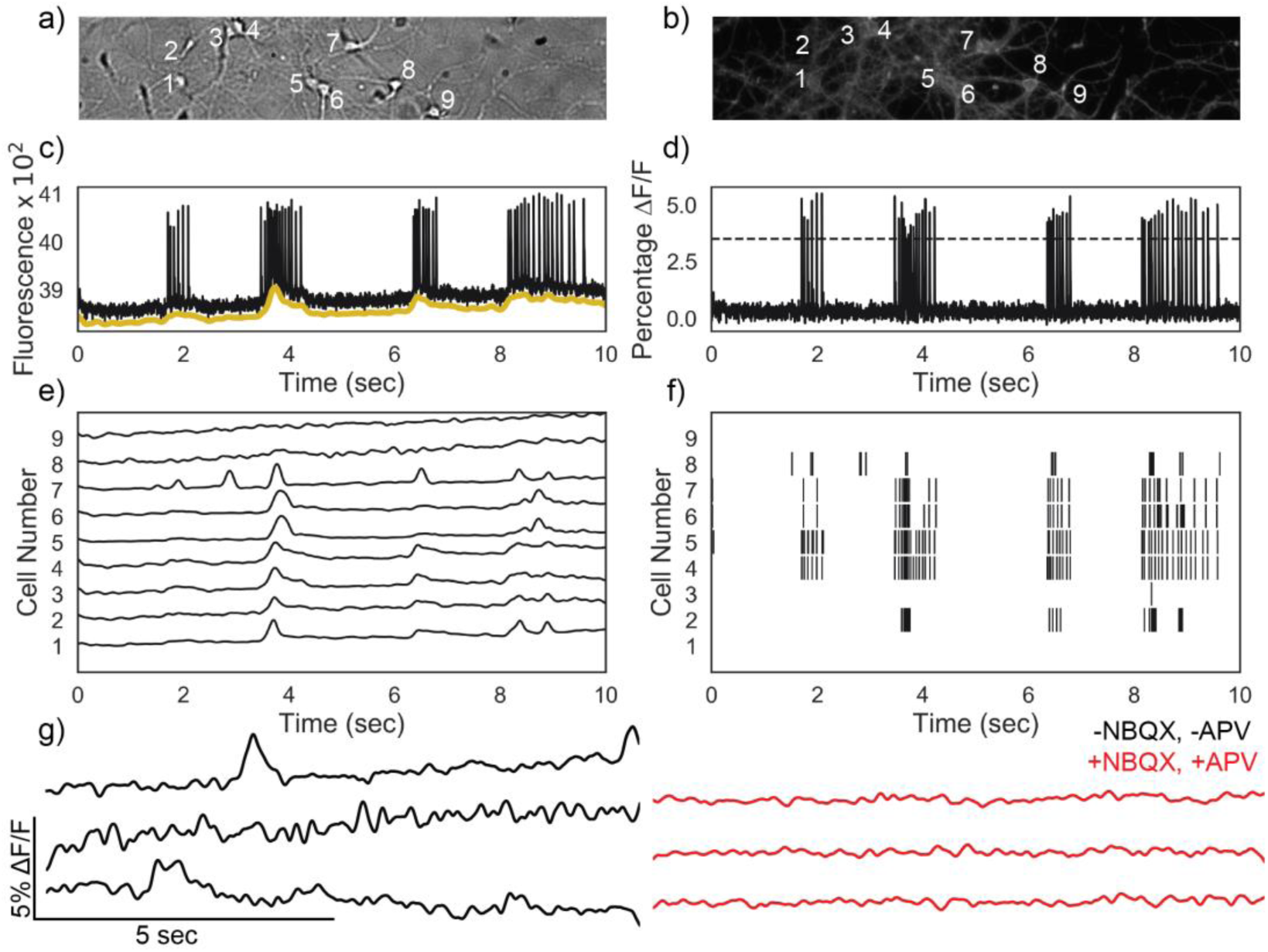
Voltage traces contain both spiking and subthreshold event data, which can be separated via regression. a, b) Brightfield and BeRST 1 fluorescence of a single field of view. c) The raw voltage imaging traces (example shown in black) were separated into “subthreshold” (gold, offset) and “spiking” traces by calculating the baseline with asymmetric least squares regression and dividing it out of the raw trace. d) The spiking trace was then converted to a spike train via thresholding based on the standard deviation of the trace and using a 2-frame rearm factor. e, f) A sample set of plots from a single field of view contains information about both spiking events and subthreshold activity. g) Variation in the subthreshold trace can be abolished with NBQX and APV.

### Using Subthreshold Activity as “Input,” Spike-Triggered Averaging Reveals Sialidase-Induced Changes in Neuronal Excitability

Spike-triggered averages (STAs) are commonly used in sensory neuroscience to reveal the “preferred inputs” of the neurons in question that elicit an action potential(Pillow and Simoncelli, 2006; Schwartz et al., 2006). STAs typically cannot be used in cultured neurons being observed with multielectrode arrays because the input into each neuron is unknown and not experimenter-controlled. By using voltage imaging and separating subthreshold and spiking activity, STAs can be generated using the subthreshold waveform as the “input” to each neuron. This allows us to measure any changes in the action potential threshold that occur due to our interventions and serves as a direct measure of neuronal excitability.

To validate the use of STA of subthreshold activity as a measure of excitability, we computed the STA of cultured mouse hippocampal neurons for a variety of interventions and culture ages. We noted that +APV, +NBQX neurons that were electrically stimulated to spike had flat STAs (d = – 1.825), p < 0.00001) relative to WT (**Supplementary Figure 4**), indicating that the majority of variation in the subthreshold traces was indeed due to subthreshold input from other neurons. We further observed that as the cultures aged, the STA height at the action potential changed minimally (6DIV vs. 12DIV: d < −0.07, p = 0.74, Permutation test, 100,000 shuffles, **Figure 3a, b**), in line with the expectation that the increased firing rate as a culture ages is primarily driven by synapse formation and less so by changes in excitability. Addition of serotonin (to increase inhibition) demonstrated that firing rate changes (−0.7 Hz) that are not driven by changes in excitability minimally affect the STA (d = 0.01, p = 0.709, Permutation test, 100,000 shuffles). Addition of strychnine (to increase bursting) to the cultures demonstrated that firing rate changes (+1.3 Hz increase, respectively) that result in synchronized bursting activity are detected by the STA (d = 1.28, p < 0.00001, Permutation test, 100,000 shuffles, **Supplementary Figures 5 and 6**).

**Figure 3.**
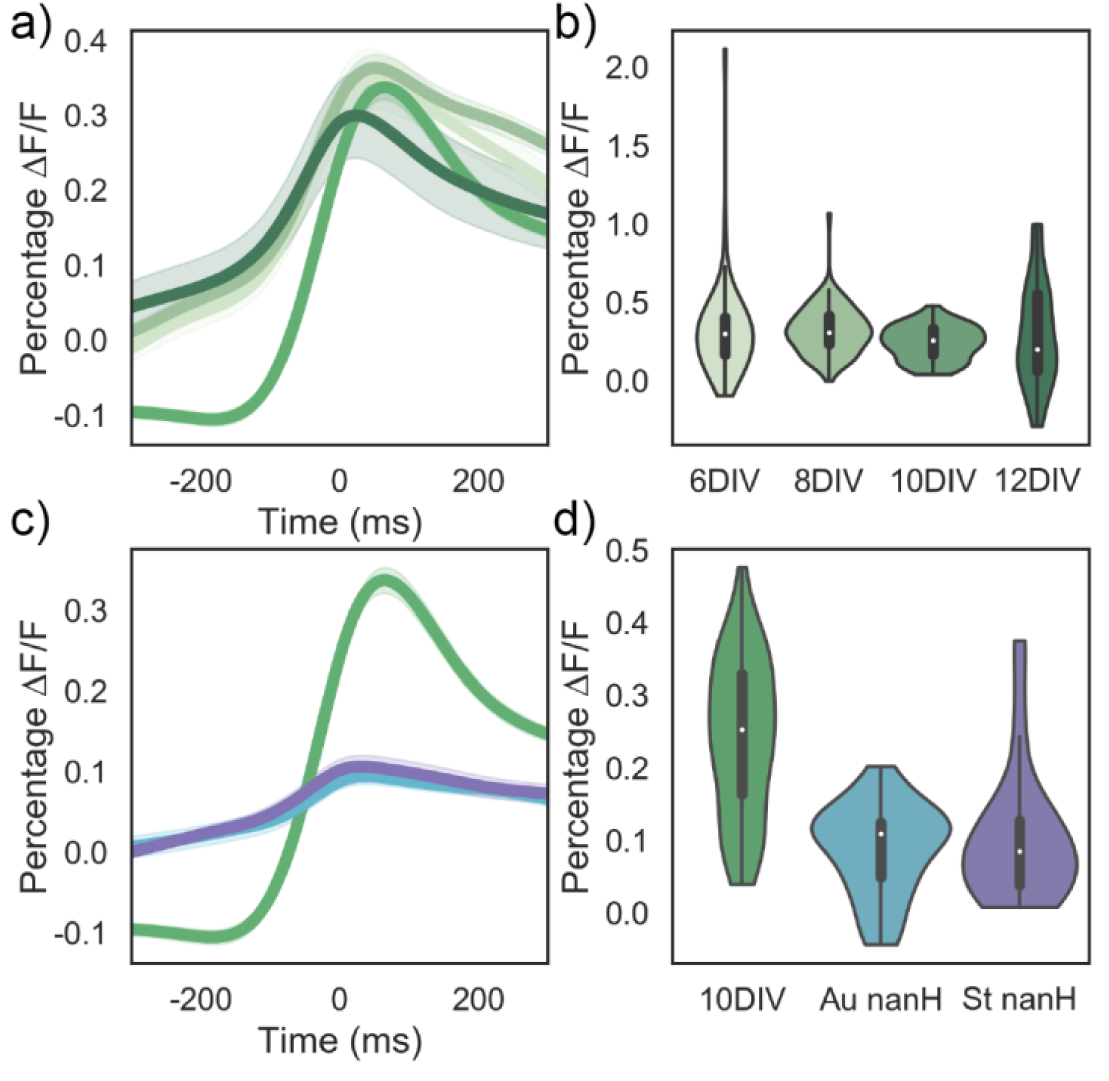
Spike-triggered averaging reveals that neuronal excitability does not markedly change as cultures age but increases in response to sialidase treatment. a) Overlay of the spike-triggered averages +/- standard deviation for 6, 8, 10, and 12DIV (in darker shades of green, n = 34, 140, 168, and 117 firing cells from 3 animals). b) At t = 0 respective to the spike, the amplitude of the STA increases slightly as the neurons age. c) Overlay of the spike-triggered averages +/- standard deviation for 10DIV, Au nanH, and St nanH (in cyan and magenta, respectively, n = 168, 44, and 66 firing cells from 3 animals). d) At t = 0 respective to the spike, both Au nanH and St nanH treatment cause a large decrease in STA amplitude. Shaded areas in a) and c) are ±SEM. Error bars in b) and d) are ±STDEV.

With this metric in hand, we next examined the STAs for 10DIV neurons treated with the nanH enzymes. The STA for neurons treated with *Salmonella* nanH indicated a large decrease in STA amplitude (d = −1.16, p < 0.00001, Permutation test, 100,000 shuffles, **Figure 3c, d**), suggesting that the observed increase in firing rate was primarily driven by a decrease in action potential threshold. This demonstrates a previously unknown role of sialic acid in the neuron and potentially explains why the brain possesses such a large amount of 2,3-linked sialic acid. The STA for neurons treated with *Arthrobacter* nanH also demonstrated a large decrease in STA amplitude (d = −1.28, p < 0.00001, Permutation test, 100,000 shuffles, **Figure 3c, d**). We noted that because a large portion of the population in the *Arthrobacter* nanH-treated samples were silent, the change in STA was primarily driven by a small proportion of the population and most cells may have undergone a decrease in excitability. To test this hypothesis, we stimulated enzyme-treated cultures with depolarizing current steps, which revealed that both treatment with *Arthrobacter* nanH and *Salmonella* nanH resulted in a significant decrease in rheobase current relative to a vehicle control (d = −1.16, p = 0.00176 and d = −1.23, p = 0.00057, respectively, **Supplementary Figure 7**). These data suggest that 2,3-linked sialosides play a role in maintenance of neuronal excitability. Because *Arthrobacter* nanH promiscuously cleaves several types of sialic acid glycosidic bonds, we reasoned 2,6- and/or 2,8-linked sialic acids may mediate a decrease in firing rate that counteracts the changes in excitability caused by cleaving 2,3-linked sialic acids.

### Synchronicity Can Be Measured Via Pairwise Cross-Correlations Between Subthreshold Traces

In order to explore other potential mechanisms by which *Arthrobacter* nanH decreased firing rate, we chose to look at network synchronicity – if the neurons were receiving significantly less input from the network, it would explain why only the most excitable cells in the network were still able to fire. A disruption in network input would result in less synchronicity between neurons. We chose pairwise cross-correlation as a metric of culture synchronicity. Specifically, we performed pairwise cross-correlation on each neuron-pair’s subthreshold trace and not on the spike trains, as is more commonly done(Tchumatchenko et al., 2011). This is because cross-correlation of spike trains tends to be highly dependent on firing rate(Dean and Dunsmuir, 2016) (**Figure 4b, e**), which would pose a problem given that the sialidase interventions significantly changed the neuronal firing rates. On the other hand, the subthreshold traces are continuous regardless of the number of spikes in each neuron and are therefore less vulnerable to artifacts that binned spike trains may cause when comparing data sets with changes in average firing rate (**Figure 4a, d**). Furthermore, using the subthreshold traces allows us to specifically compare the input patterns of each neuron pair, whereas cross correlating the spike trains might conflate synaptic input with the excitability filter intrinsic to each neuron that affects spiking probability.

**Figure 4.**
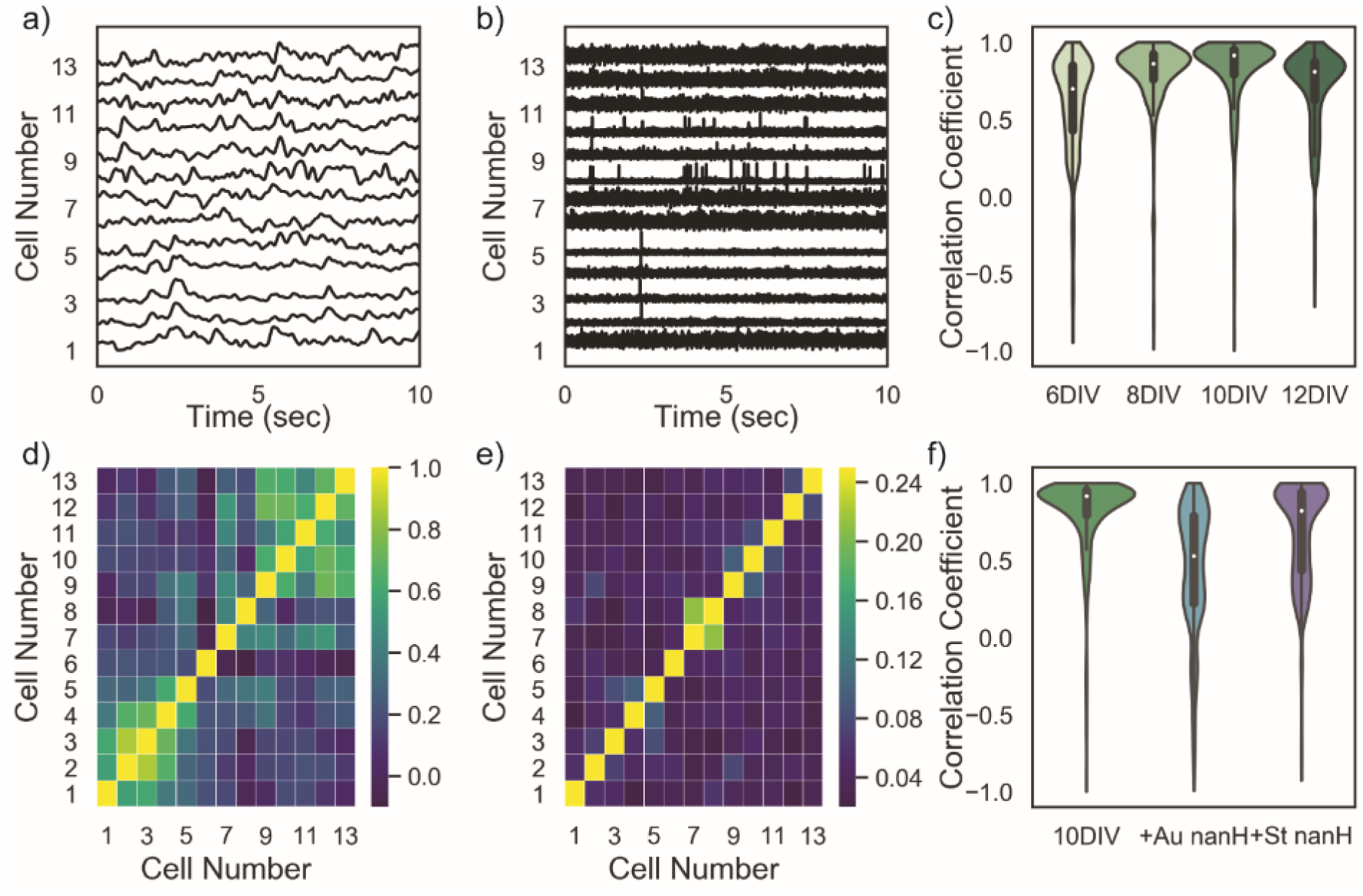
Pairwise cross-correlation of the subthreshold traces reveals the degree of similarity between synaptic inputs to measured neurons. a, b) Cross correlation could be performed on either the subthreshold trace (a) or the spike trace (b), resulting in the heatmaps d) and e), respectively. c) As the cultures aged, the average normalized cross-correlation value approached 1 before day 12 (n = 76, 144, 191, and 125 cells across 3 animals). f) Treatment with Au nanH resulted in a large decrease in average cross-correlation, while treatment with St nanH resulted in a smaller decrease in average cross-correlations (n = 191, 108, and 98 cells per treatment across 3 animals). Error bars are ±STDEV.

As expected, we observed an increase in pairwise cross-correlation as the neurons aged from day 6 to day 12 (d = 0.31, p < 0.00001, Permutation test, 100,000 shuffles, **Figure 4c**), concordant with an increase in culture connectivity as the neurons mature and form synapses. Treatment with APV and NBQX resulted in a large decrease in pairwise cross-correlation values as compared to untreated cells (d = −1.7, p < 0.00001, Permutation test, 100,000 shuffles, **Supplementary Figure 7**), consistent with the hypothesis that subthreshold traces reflect summed synaptic input and that traces with high correlations are experiencing similar inputs. Treatment with *Salmonella* nanH resulted in a small decrease in average pairwise cross-correlations (d = −0.47, p < 0.00001, Permutation test, 100,000 shuffles, **Figure 4f**), suggesting that 2,3-linked sialic acids do not play a large role in neuronal connectivity. Treatment with *Arthrobacter* nanH resulted in a large decrease in average pairwise cross-correlations (d = −0.98, p < 0.00001, Permutation test, 100,000 shuffles, **Figure 4f**), indicating that the cultures had undergone a large decrease in synchronicity and potentially explaining the decrease in overall firing rate despite the concomitant increase in excitability.

Both nanH treatments resulted in bimodal distributions of pairwise cross-correlations, potentially suggesting that a large group of cells became disconnected from the greater network. Another potential explanation is that the enzyme treatment resulted in the initially synchronous network activity breaking up into multiple “islands” of neuronal networks. Synchronicity on its own is not able to tease apart these potential mechanisms, so we sought other measures of network activity.

### Dimensionality Reduction Models Neuronal Network Connectivity

We reasoned that the distinction between some neurons no longer being connected to the network and the formation of several smaller networks (**Figure 5a, b**) is a question of “network integration” – that is, the degree to which a neuron’s activity is being driven by the network at large. In order to model network integration, we applied factor analysis, a well-known dimensionality reduction method, to the subthreshold data.

**Figure 5.**
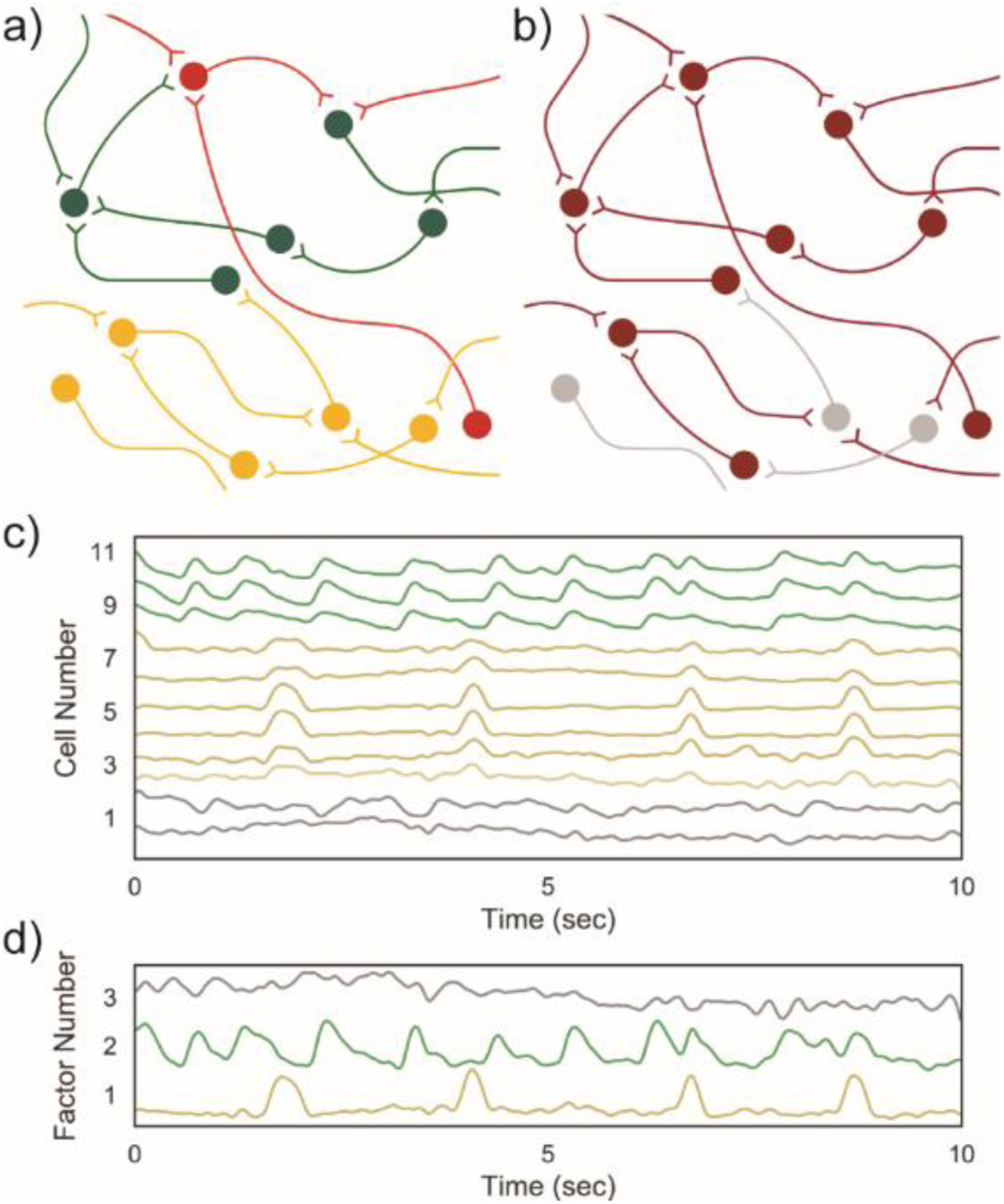
Factor analysis of subthreshold traces models network inputs that explain the subthreshold variation in the measured neurons. a, b) Reduced correlation between neurons in a single field of view may be driven by the neurons forming “islands” of similar activity (a) or some neurons losing their connection to the greater network (b). c) The activity of 11 neurons in a single field of view can be modeled as the result of a small number of “hidden variables” that represent network-driven activity. d) Fitting factor analysis with three dimensions reveals that 50% of the variance in network activity is explained by factor 1 (bottom), with factor 2 explaining another 20% and factor 3 explaining 8% of remaining variance, respectively.

Dimensionality reduction techniques are commonly-used in neuroscience for simplifying visualization and analysis of high-dimensional data sets(Pang et al., 2016). Principal Component Analysis (PCA) is perhaps the most commonly used dimensionality reduction technique, having been used to analyze variance in action potential waveforms, neural population dynamics, and identifying ensembles in a large population of cells(Aljadeff et al., 2016). PCA, however, seeks to maximize explained variance in each component, and is therefore most suited for modeling properties of the measured neurons that manifest as sources of increased variation. Shared synaptic input due to network activity, however, should manifest as correlations, or covariance, between recordings of several neurons that are measured simultaneously. PCA does not aim to preserve covariance between measurements and is therefore inappropriate for modeling network activity. Factor analysis (FA) is a related dimensionality reduction technique that seeks to maximize the covariance between measured neurons when constructing factors(Athalye et al., 2017; Soletta et al., 2017). In other words, FA can be used to model the “hidden” network state that is driving the activity of the “visible” measured neurons by assuming that correlations between the activities of the measured neurons are being driven by similar inputs (**Figure 5c, d**).

FA separates each measured neuron’s activity into a “shared variance” term and a noise, or “private variance,” term – that is, the technique assumes that some portion of the measured activity in each neuron is driven by inputs shared with other measured neurons, while some portion is driven by inputs unique to that neuron or neuron-specific spontaneous activity(Athalye et al., 2017). By labeling neurons with the percentage of their variance that is shared covariance, factor analysis provides us with a metric that models the “connectedness” of each neuron to the network at large. Given that this provides a single measure per neuron, we believe it is uniquely suited for assaying the overall connectivity of a network on a per-neuron basis. Much like PCA, the number of factors generated by FA must be chosen beforehand and the eigenvalues of the factors generated by FA can be visualized on a scree plot to determine the appropriate number of factors. The scree plot analysis can be used, in our experience, to assay the maturity of the network – at 10DIV, when the network is mature, a vast majority of the observed variance can be explained by a single factor, while at 6DIV, multiple factors are required to explain a majority of the observed variance (**Supplementary Figure 9**). In general, we posit that factor analysis provides a much more direct method to assay network integration and can be used in conjunction with synchronicity to describe network properties in cultured neurons.

### *Arthobacter* Sialidase Treatment Reduces Neuronal Shared Variance

We began by performing FA with three potential factors on each field of view in the time course data set from 6 to 12 DIV. As the cultures aged, the per-neuron shared variance proportions were high and increased negligibly (+9% from 6DIV to 10DIV, p = 0.02028, Permutation test, 100,000 shuffles, **Figure 6a**) and the number of factors required to explain most of the variance decreased (50% of fields multidimensional at 6DIV to 10% of fields multidimensional at 10DIV). These results suggest that in culture, the majority of neuronal membrane potential variation is being driven by the network even at early timepoints, but immature cultures have not formed a large number of connections, resulting in several “islands” of neurons that have formed small networks. This hypothesis was also supported by the observations that the total network variance explained by all three factors did not significantly change from 6DIV to 10DIV (+4% change, p = 0.48159, **Figure 6d**).

**Figure 6.**
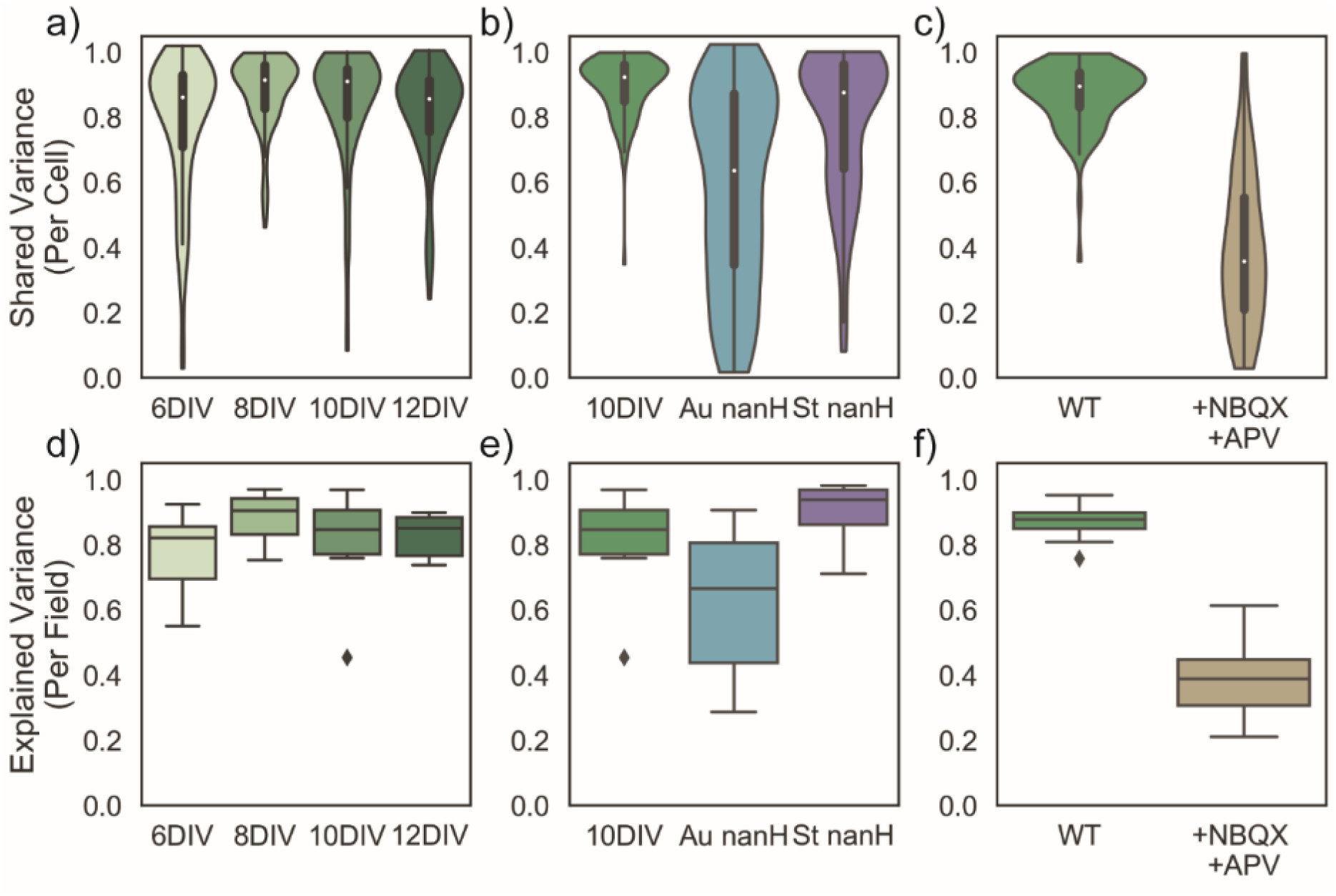
*Arthrobacter* sialidase treatment results in a large decrease in shared variance, suggesting that a large number of neurons are no longer well-connected to the network. a, d) As the cultures age, they experience negligible changes in the proportion of their variance that is explained by network activity (n = 76, 144, 191, and 125 cells across 3 animals). b, e) Treatment with Au nanH results in a large decrease in neuronal shared variance and the proportion of observed variance explained with 3 factors, while treatment with St nanH results in negligible change in both types of variance (n = 191, 108, and 98 cells per treatment across 3 animals). c, f) Treatment with NBQX and APV results in a large decrease in shared variance per neuron and proportional explained variance per field of view. Outliers are marked with black diamonds and n = 89 and 58 cells across 10 fields of view for each sample. Error bars are ±STDEV.

To verify that factor analysis measures network integration, we performed factor analysis on networks treated with NBQX and APV, and found a large decrease in per-neuron shared variance (−47% change in shared variance, p < 0.00001, Permutation test, 100,000 shuffles) and in variance explained by three factors (−46% change in explained variance, p < 0.00001, Permutation test, 100,000 shuffles, **Figure 6c, f**) suggesting that the integration of the measured neurons into a network had been significantly disrupted. Further analysis of interventions that modulate firing rate without interfering with synaptic transmission, such as serotonin, revealed small changes in per-neuron shared variance after drug application (−6% change in shared variance, p = 0.93645, Permutation test, 100,000 shuffles, **Supplementary Figures 5**). Meanwhile, strychnine application, which results in synchronized bursting, causes a small increase in shared variance and a large reduction in the spread of shared variance (+4% increase, p = 0.00003, Permutation test, 100,000 shuffles, **Supplementary Figures 6**). We reasoned that, while performing factor analysis on subthreshold traces is much more resistant to spike frequency-related artifacts than performing the analysis on spike trains, large changes in the spiking frequency across the entire coverslip would result in changes to the amount of excitatory input each neuron receives. Because the neurons act spontaneously during lulls in excitatory input, cultures experiencing more excitatory input will have higher per-neuron shared variance even if connectivity is unchanged. With these controls in hand, we then performed factor analysis on networks treated with nanH enzymes.

Upon treatment with *Salmonella* nanH, FA revealed a small reduction (−6% change, p < 0.00001, Permutation test, 100,000 shuffles) in neuronal shared variance (**Figure 6b, e**), suggesting that the increase in firing rate was indeed driven by a change in excitability and little change in network integration. Treatment with *Arthrobacter* nanH, however, resulted in a large reduction in network shared variance (−26% change, p < 0.00001, Permutation test, 100,000 shuffles, **Figure 6b, e**), which is the likely main driver of the resulting decrease in overall firing rate. Using factor analysis made it clear that the bimodal distribution of pairwise cross-correlations in *Arthrobacter* nanH treated-neurons was caused by a large number of neurons becoming disconnected from the network. This is in contrast to 6DIV untreated neurons, whose bimodal distribution of pairwise cross-correlations was a result of smaller networks of neurons. Given that neuronal cell surface 2,6-linked sialic acids are primarily attached to proteins, which include ion channels and receptors(Schnaar et al., 2014), this analysis suggests that charged sialic acids attached to these proteins play an important role in efficacy of synaptic transmission.

## Discussion

Sialic acids play a fundamental role in many neuronal processes, such as development and plasticity(Schnaar et al., 2014; Tong et al., 2017). Gross removal of sialic acid residues has also been demonstrated to decrease spiking activity in neuronal networks(Isaev et al., 2007). Here we developed a voltage imaging platform to show that 2,3-linked sialic acid residues modulate excitability in mouse hippocampal networks, while 2,6- and 2,8-linked sialic acid residues may play a role in network integration.

We observed that firing rate of cultured hippocampal neurons was greatly decreased by global removal of cell-surface sialic acid, in concordance with the observations in hippocampal slices(Isaev et al., 2007). Global desialylation resulted in a large decrease in network integration, resulting in a loss of synchronized synaptic input to a large number of neurons within each network and therefore a decrease in observed firing rate. This suggests that 2,6- and 2,8-linked sialic acid residues have a localized influence on transmission at the synapse, perhaps by modulating the conductance or binding properties of the receptors that they are linked to.

In contrast, selective removal of just the 2,3-linked sialosides resulted in an increase in action potential firing rate driven by a decrease in action potential threshold. Furthermore, loss of 2,3-sialosides did not significantly affect network integration, suggesting that these glycans, which are predominantly on neuronal gangliosides(Schnaar et al., 2014), play a very different role in neuronal electrophysiology than 2,6- or 2,8-linked sialic acids. We speculate that the neuron uses the sialylation state of different populations of membrane biomolecules, such as gangliosides, ion channels, and receptors, to finely modulate their electrophysiological behavior. This is consistent with the presence of multiple endogenous neuraminidase enzymes in the brain that possess different substrate specificity and are actively regulated in response to electrophysiological activity(Pshezhetsky and Ashmarina, 2018). Prior studies demonstrating that blocking endogenous neuraminidase activity affects network spiking properties further support this hypothesis(Isaeva et al., 2010).

Because voltage imaging is analogous to intracellular electrophysiology in the sense that directly measures voltage across the cell membrane, we were able to extract the subthreshold activity from the raw voltage trace for further analysis. Many network analyses that are traditionally performed on spike trains generated by extracellular recording, such as pairwise cross-correlation and dimensionality reduction, are easily ported to the subthreshold traces and reveal similar information about the neuronal networks. However, using the subthreshold traces avoids some of the spiking frequency-induced artifacts associated with preparing spike-train data for input into these methods. The subthreshold traces can also be used to generate spike-triggered averages that interrogate the action potential threshold on a per-neuron basis.

In conclusion, we find several pieces of evidence to suggest that different populations of sialic acid play differential roles in network electrophysiology: 1) Removal of cell surface sialic acid with a pan-sialidase resulted in a decrease in firing rate that 2) was attributed to a decrease amount of network-driven input to the observed neurons; 3) Removal of cell surface sialic acid with a 2,3-sialidase resulted in an increase in firing rate that 4) was attributed to an increase in excitability via a decrease in input required to reach the action potential threshold in the observed neurons. These findings show that changes in glycosylation state alone can drive changes in neuronal electrophysiology. In light of this result, it is now clear that our understanding of neural communication cannot be complete without an understanding of the currently-underexamined neuronal glycosylation state. We further note that several of these analyses were made possible by voltage imaging: 1) High-throughput measurement of neuronal spiking generated a large dataset that 2) contained both spiking and subthreshold information, enabling observational measurement of neuronal excitability; 3) Simultaneous measurement of several neurons facilitated measurement of network properties that are difficult to measure with traditional intracellular electrophysiology, such as synchronicity and network integration. We believe this platform for studying neuronal network properties in culture will prove useful for modeling any disorder or dysfunction that results in network-level changes to electrophysiology.

## Acknowledgements

We would like to thank Katherine Derosier for helpful discussions regarding statistical modeling. We would also like to thank Melissa Anne Gray and Lisa Nakayama for expert technical support with sialidase enzymes and neuronal culture, respectively. We also thank Gloria Ortiz and Evan W. Miller for providing BeRST 1. This work was generously supported by the US National Institutes of Health (CA200423).

## Author Contributions

Conceptualization, RUK and CRB.; Methodology and Formal Analysis, RUK., Investigation, RUK and CLW.; Writing – Original Draft, RUK and CRB.; Writing – Review and Editing, CLW and CRB.; Funding Acquisition – CRB.; Supervision, CRB.

## Methods

### Dissociated hippocampal cultures

All animal procedures were approved by Stanford University’s Administrative Panel on Laboratory Animal Care and conformed to the NIH Guide for Care and Use of Laboratory Animals and the Public Health Policy. Primary hippocampal tissue was harvested from E16.5 C57BL/6 embryonic mice (Charles River) immediately after sacrifice of the pregnant dam. The isolated hippocampi were dissociated using Papain Dissociation System (Worthington Biochemical Corporation) and trituration with fire-polished Pasteur pipettes. The dissociated cells were plated onto 12mm coverslips (Chemglass Life Sciences) pre-treated with Poly-D Lysine (PDL; 1 mg/mL, Sigma-Aldrich) at a density of 6 ×s 10^4^ cells per coverslip. The cells were maintained for 24 hours in Dulbecco’s modified eagle medium (DMEM; Gibco) supplemented with 4.5 g/L D-glucose, 10% FBS, 1% GlutaMAX, and 2% B-27 (Gibco), and then switched to Neurobasal media (Gibco) with 1% GlutaMAX and 2% B-27 supplement. Plates were incubated at 5% CO_2_, 37°C, and subsequent media changes took place every 7 days.

### Voltage imaging of neurons

Neurons were incubated with an HBSS solution (Gibco) containing BeRST1^12^ (1.0µM; Miller Lab, UC Berkeley) for 20 minutes at 37°C prior to imaging. Functional imaging of hippocampal cells was performed on an Axio Observer Z-1 (Zeiss) equipped with a Spectra-X Light engine LED light (Lumencor), controlled with Micro-Manager. Images were acquired with a Plan-Apochromat 20x/0.8 air objective (Zeiss) paired with image capture from the ORCA-Flash4.0 sCMOS camera (sCMOS; Hamamatsu) with 4×4 in-camera binning. We used a sampling rate of 0.5 kHz over 5 or 10 seconds in order to resolve individual action potentials. To achieve maximum sampling rate, a field of view (FOV) size of 512×100 pixels (665.6×130 µm) was used for simultaneous recording of 5-15 neurons at a time. BeRST 1 was excited with a 631 nm light (LED; 631nm, 28 nm bandpass) with an LED power of 70%. Emission from BeRST 1 was collected with a 680/10 nm bandpass emission filter after passing through a dichroic mirror (425/30 nm, 514/30 nm, 592/25 nm, 709/100 nm LP).

To assess network maturation, neurons were imaged on 6, 8, 10, and 12 DIV. To investigate the effects of bond-specific sialic acid removal, 10 DIV neurons were incubated with *Salmonella typhimurium* nanH (St nanH; 2 μM) and *Arthrobacter ureafaciens* nanH (Au nanH, 2 μM) in Neurobasal media at 37°C for 1 hour prior to imaging. Enzymes were prepared as described in Gray, 2019(Gray et al., 2019). Enzyme selectivity was validated via lectin staining using fluorescein-labeled *Sambucus nigra* lectin (SNA; Vector Laboratories) and biotinylated *Maackia amurensis* Lectin II (MAL-II; Vector Laboratories). Cells were fixed with 4% PFA (15 min), blocked with 2% goat serum (1 hr), and then incubated with SNA (10μg/mL) and MAL-II (10μg/mL) for one hour at room temperature. MAL-II imaging was done by conjugation with streptavidin functionalized with Alexa 594 (Vector Laboratories).

In experiments involving pharmacologic validation of neuronal network properties, subthreshold activity was abolished with glutamate receptor antagonists 2,3-Dioxo-6-nitro-1,2,3,4-tetrahydrobenzo-[f]quinoxaline-7-sulfonamide (NBQX; 10 μM; Santa Cruz Biotechnology) and DL-2-Amino-5-phosphonopentanoic acid (APV; 50 μM; Sigma-Aldrich). Neurons were also treated with serotonin (5 μM; Sigma-Aldrich) and strychnine (5 μM; Sigma-Aldrich) to assess relative levels of network excitation and inhibition. All treatments were applied to neurons via supplementation to the HBSS imaging solution.

Extracellular field stimulation was delivered by a SIU-102B bipolar stimulator (Warner) with triggering provided through an Isolated Pulse Stimulator (Model 2100; A-M Systems) via TTL cable. Action potentials were triggered 10-ms 5-30 mA bipolar current injections or 80 V voltage clamps delivered at 2 Hz. To prevent recurrent activity, the HBSS bath solution was supplemented with 10 μM NBQX and 50 μM APV as described above.

### Image analysis

Analysis of voltage traces in primary neurons was performed using ImageJ and custom Python scripts. All Python scripts used to analyze data in this study are available at github.com/rishi-kulkarni/SpykeMapper. Regions of interest (ROIs) were drawn around cell bodies within a field of view and the average fluorescence over time was extracted and inputted into an Excel workbook. The fluorescence time course data was then analyzed using a custom Python script that performed subthreshold trace extraction and spike train generation. Briefly, the subthreshold activity was identified using asymmetric least squares regression and subtracted from raw time course data to generate a flattened trace containing a flat baseline and spiking activity. Spikes were identified from the flattened trace using a threshold of +6 STDEV of all cellular fluorescence values within a coverslip to generate a digitized spike train containing all-or-nothing firing data or Raster plot. We quantified the amount of neuronal input necessary to produce spiking activity under different conditions using the spike-triggered average (STA) as a metric for excitability. Spike-triggered averages for each cell were generated by summing baseline values +/- 300 ms around each spike and dividing by the total number of spikes. The STA amplitudes were compared using Cohen’s d. Cross-correlations were calculated using the numpy.correlate function with a maximum time-lag of 100 ms. The average pairwise cross-correlation values per network were compared using Cohen’s d. Factor analysis was performed using the FactoryAnalyzer module with no rotation. The shared variance values per network were compared using Cohen’s d.

### Hypothesis testing

All statistical hypothesis testing was performed via permutation testing by testing for changes in mean with 100,000 permutations.

