## Supplemental Figures for "Subthreshold Voltage Analysis Demonstrates Neuronal Cell-Surface Sialic Acids Modulate Excitability and Network Integration"

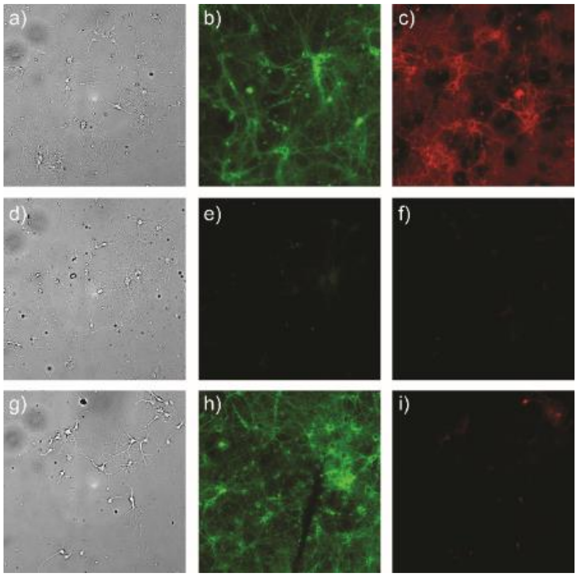


**Figure S1.** Mouse hippocampal neurons (a, d, g) were treated with a heat-inactivated *Arthrobacter* nanH (a, b, c), active *Arthrobacter* nanH (d, e, f), and active *Salmonella* nanH (g, h, i). Lectin staining with SNA (b, e, h) and MAL-II (c, f, i) reveals that *Arthrobacter* nanH cleaves both 2,6- and 2,3-linked sialic acids, while *Salmonella* nanH prefers 2,3-linked sialic acids.

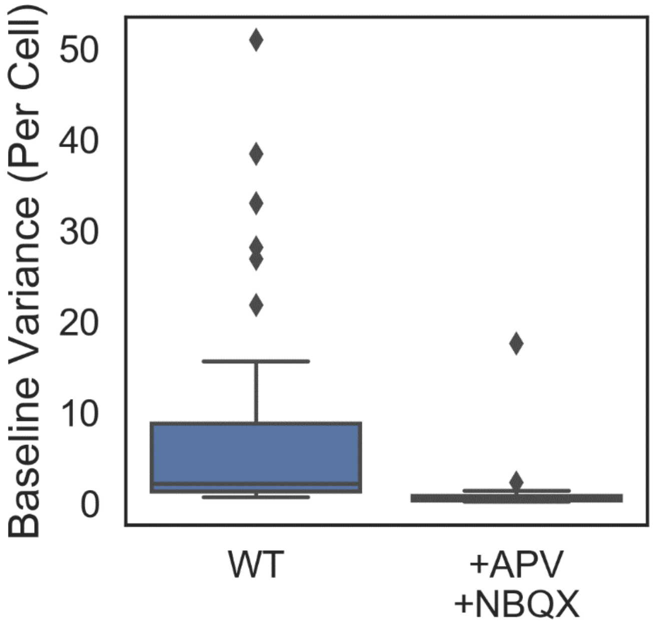


**Figure S2.** Addition of APV and NBQX to WT mouse hippocampal neurons results in a large reduction of variance of the calculated baseline, suggesting that the baseline reflects summed synaptic activity. n = 53 and 34 cells, respectively. p = 0.00059 and d = -0.701.

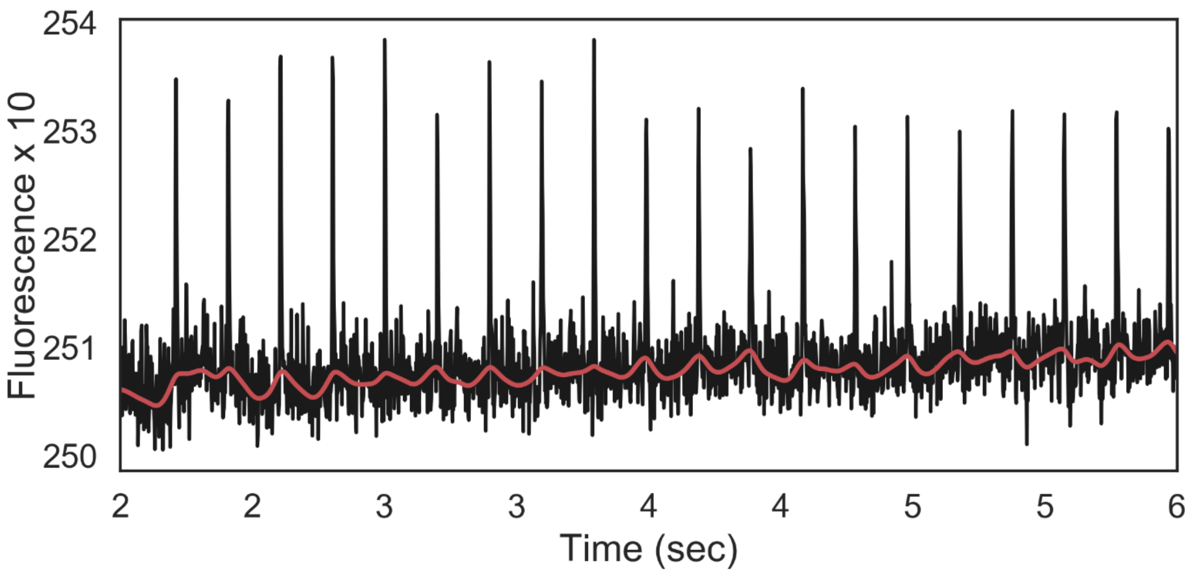


**Figure S3**. Inducing action potentials via field stimulation in mouse hippocampal neurons treated with APV and NBQX results in a relatively flat, low-variance subthreshold fit, suggesting that the major source of variance in the fit is not due to artifacts caused by dye-spike interactions.

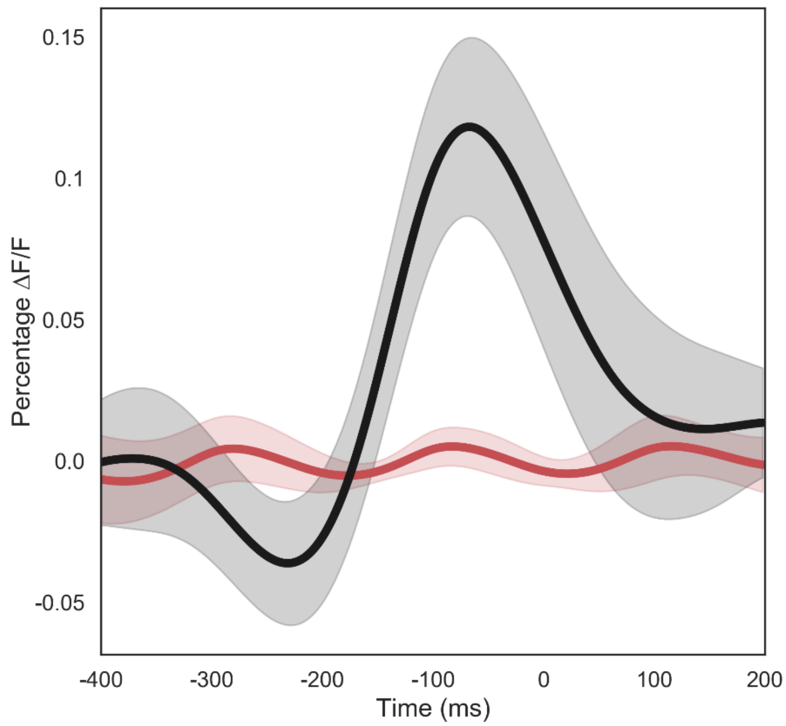


**Figure S4**. Addition of NBQX and APV to cells that were then stimulated to fire action potentials via field stimulation (red) resulted in a flat STA relative to the large changes in amplitude seen in the same cells firing spontaneously (black). $n = 7$ cells for each sample, $p < 0.00001$, $d = 1.825$. Shaded areas represent ±STDEV.

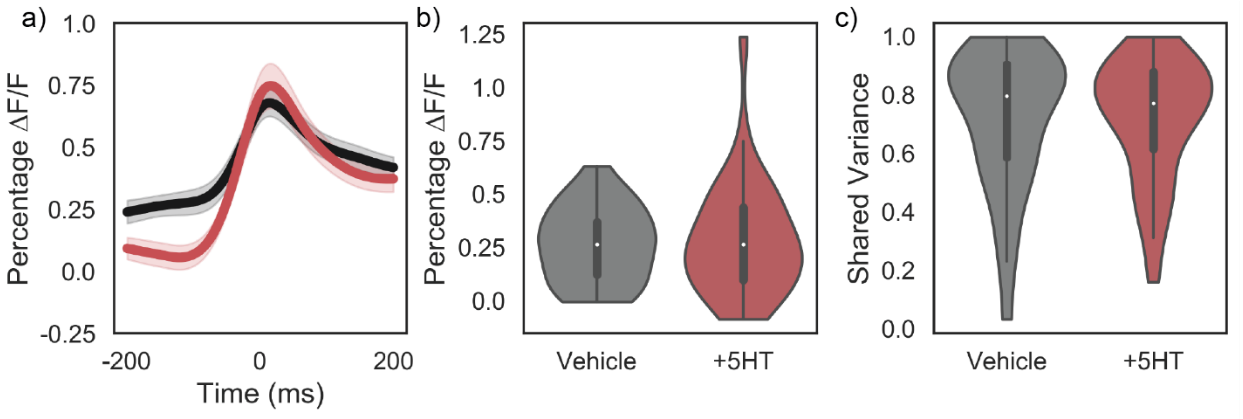


**Figure S5.** a, b) Addition of serotonin to imaging media results in a negligible change in STA amplitude at $t$ = 0 (p = 0.709, d = 0.10). Shaded areas represent ±STDEV. c) Per-neuron shared variance is minimally affected by serotonin treatment (p = 0.93645, d = 0.01). n = 43 and 58 cells from 3 coverslips.

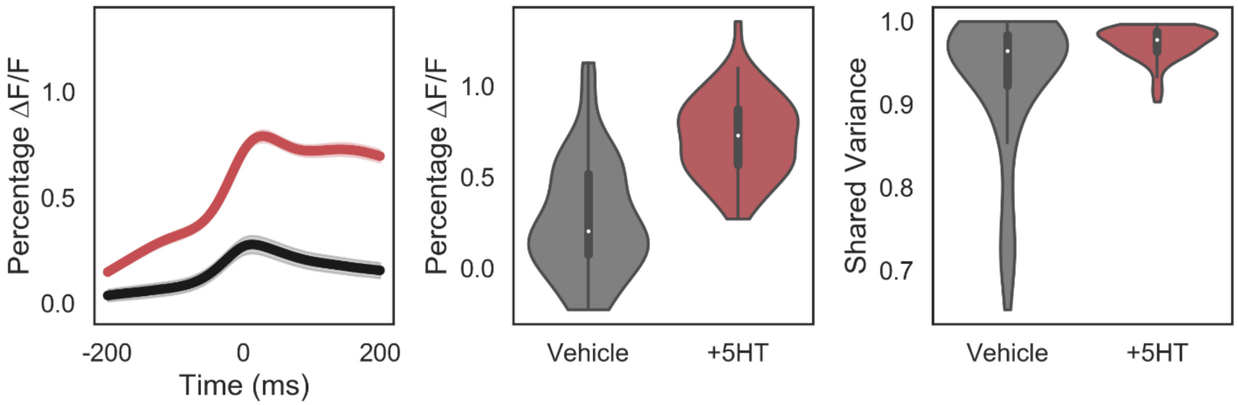


**Figure S6.** a, b) Addition of strychnine to imaging media results in an increase in STA amplitude due to increased bursting ($p < 0.00001$, $d = 1.28$). Shaded areas represent ±SEM. c) Per-neuron shared variance is increased by strychnine treatment ($p = 0.00005$, $d = 0.66$) $n = 51$ and 46 cells from 3 coverslips.

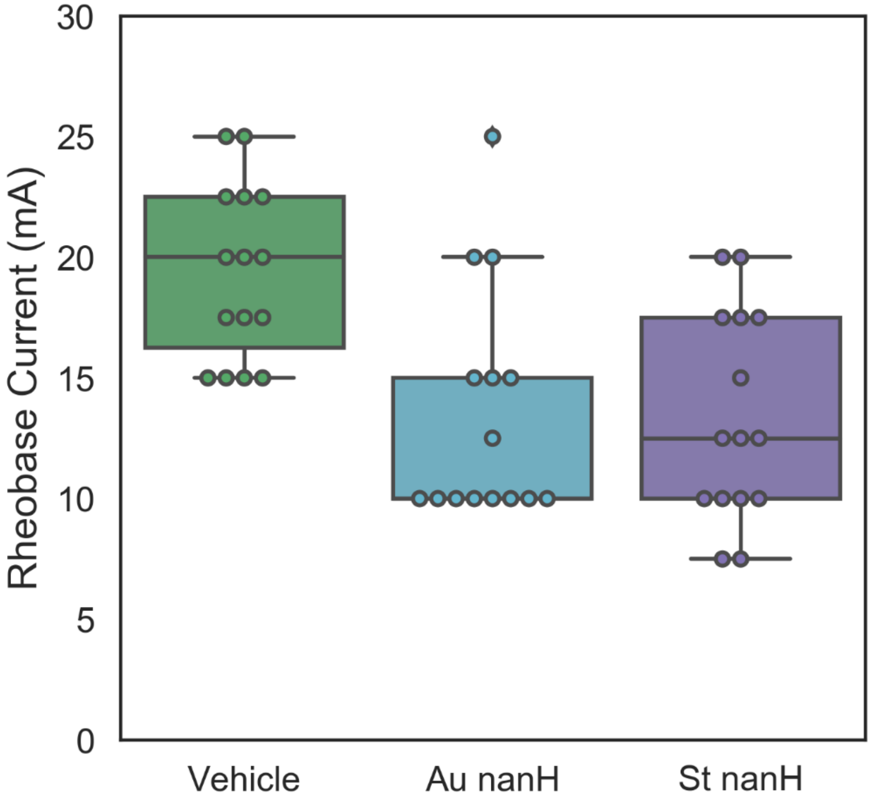


**Figure S7.** Rheobase current measurement with extracellular field stimulation. Error bars are ±STDEV for n = 15 cells per sample.

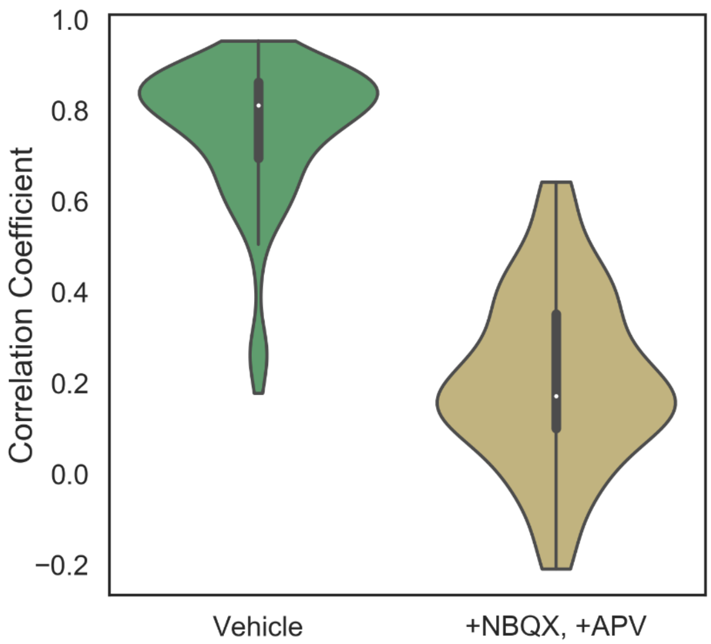


**Figure S8.** Addition of NBQX and APV to wild type mouse hippocampal neurons results in a large reduction in pairwise cross-correlation values between subthreshold traces ($\Delta = -1.7$, $p < 0.00001$, n = 42 and 31 cells across 3 coverslips).

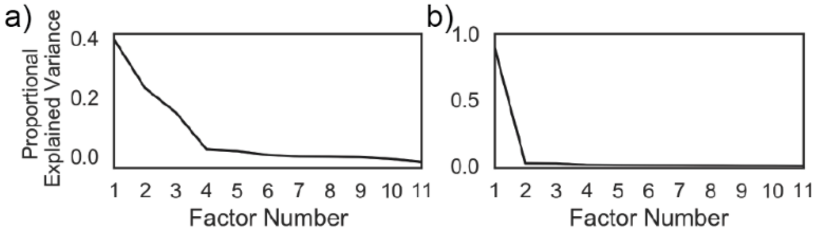


**Figure S9.** 6DIV neurons (a) require a larger number of factors to explain the amount of variance explained by a single factor in 10DIV neurons (b).
